# Inhibition of the type IV secretion system from antibiotic-resistant *Helicobacter pylori* clinical isolates supports the potential of Cagα as an anti-virulence target

**DOI:** 10.1101/2023.09.01.555934

**Authors:** Flore Oudouhou, Claire Morin, Mickael Bouin, Christiane Gaudreau, Christian Baron

## Abstract

*Helicobacter pylori* resistance to antibiotics is a growing problem and it increasingly leads to treatment failure. While the bacterium is present worldwide, the severity of the clinical outcomes is highly dependent on the geographical origin and genetic characteristics of the strains. One of the major virulence factors identified in *H. pylori* is the *cag* pathogenicity island (*cag*PAI), which encodes a type IV secretion used to translocate effectors into human cells. Here, we investigated the genetic variability of the *cag*PAI among 13 antibiotic-resistant *H. pylori* strains that were isolated from patient biopsies in Québec. Seven of the clinical strains carried the *cag*PAI, but only four could be readily cultivated under laboratory conditions. We observed variability of the sequences of CagA and CagL proteins that are encoded by the *cag*PAI. All clinical isolates induce interleukin-8 secretion and morphological changes upon co-incubation with epithelial gastric cancer cells and two of them produce extracellular T4SS pili. Finally, we demonstrate that molecule 1G2, a small molecule inhibitor of the Cagα protein from the model strain *H. pylori* 26695, reduces interleukin-8 secretion in one of the clinical isolates. Co-incubation with 1G2 also inhibits the assembly of T4SS pili, suggesting a mechanism for its action on T4SS function.

## Introduction

*Helicobacter pylori* is a human pathogen that colonizes the stomach of about half of the world’s population. While the infection usually leads to asymptomatic gastritis, more severe outcomes such as peptic ulceration and adenocarcinoma are observed in a subset of subjects (1,2). Different treatment regimens have been established to eradicate the infection, but due to increasing resistance to antibiotics, failure of *H. pylori* eradication therapy is becoming a growing problem (3). The most virulent *H. pylori* strains carry the *cag* pathogenicity island (*cag*PAI), a 40 kb chromosomal region, and have potential to cause severe gastric diseases (4–6). The *cag*PAI comprises 28-30 genes and encodes a type IV-secretion system (T4SS), a macromolecular complex that spans across the double membrane of the bacterium enabling the injection of effectors directly into host cells. Several molecules are translocated through the *H. pylori* T4SS such as the cytotoxin CagA, bacterial peptidoglycan, chromosomal DNA and heptose-1,7-bisphosphate (HBP), a metabolic precursor of lipopolysaccharide synthesis (7–10). While all these effectors are known to contribute to pro-inflammatory responses in epithelial gastric cells via activation of the NF-κβ transcription factor, CagA is recognized as the major virulence factor.

CagA is a 120-145 kDa protein encoded by the *cag*PAI that is associated with increased risk of gastric cancer and is considered an oncoprotein (4,6). Following bacterial attachment and CagA translocation into epithelial cells, CagA can interact with more than 25 host cell proteins, and it hijacks cellular signaling pathways. CagA induces actin-based cytoskeletal rearrangements, cell motility and proliferation, it affects cell polarity, cell adhesion, the cell cycle, the apoptosis pathway and it also induces inflammatory pathways (11). The interaction of CagA with some of its partners depends on its phosphorylation status. Following its translocation, CagA undergoes phosphorylation by Src and Ab1 kinases on tyrosine sites of the EPIYA (Glu-Pro-Ile-Tyr-Ala) motifs that are present in its C-terminus (12–15). Based on the flanking sequences, four specific EPIYA motifs have been described. The EPIYA-A, -B and -C motifs are usually found in *H. pylori* isolates from Western countries while EPIYA-A, -B and -D are present in East Asian strains (16). The EPIYA-C site occurs in different numbers; it is typically present in one to three copies and these differences are directly related to the levels of CagA tyrosine phosphorylation and binding to host proteins like SHP-2 (Src homology 2 phosphatase) (16). A larger number of EPIYA-C motifs and the presence of EPIYA-D are associated with increased virulence and carcinogenicity (12). In strains from Western countries it was observed that CagA can carry a specific A/T polymorphism, called the EPIYT motif, which presents an alternative binding site for phosphatidylinositol 3-kinase (PI3-kinase), influencing cancer risk (17). In addition, the C-terminal domain includes another repeated sequence motif that was originally designated as the CagA multimerization (CM) motif enabling multimerization via this 16 amino-acids region (18). Differences between five typical amino-acid residues enable the distinction between Western and East Asian CagA CM motifs. Whereas Western species contain multiple CM motifs located within each EPIYA-C region plus one distal to the last EPIYA-C, East Asian CagA proteins comprise a single CM motif that is located distally to the EPIYA-D segment.

The injection of effectors into the host cell cytoplasm requires bacterial adhesion to the gastric epithelium. CagL that locates at the tip of pilus-like structures (T4SS pili) is one of the *cag*PAI-encoded proteins that contribute to this process. CagL binds to many host integrins, such as the integrins α5β1, αVβ3, αVβ5, αVβ6 and αVβ8, notably via the conserved RGD (Arg-Gly-Asp) motif (19–21). Its binding to host integrins can activate the NF-κβ pathway leading to the production of several pro-inflammatory cytokines. It has been shown that several variations of the CagL sequence are associated with higher risk of gastric cancer, e.g. the CagL-Y58E59 variant induces higher integrin α5β1 expression levels and increases inflammation (22,23). Several other variations at different amino acid residues (Thr30; Asn101; Ala141; Glu142; Asn201; Ile234) are proposed to occur in higher rates in gastric cancer patients, but the mechanism on their contribution requires further investigation (23).

In *H. pylori*, T4SS pilus biogenesis and translocation of effectors are energized by three ATPases (24–27). Among them, Cagα is a VirB11-like ATPase. VirB11-like proteins are present in all T4SSs and they have been extensively characterized (28–32). Because of its importance for T4SS functions, Cagα is a target for T4SS inhibitors. We have previously identified the small molecule 1G2 that binds Cagα and reduces its ATPase activity *in vitro* (33). We also showed that 1G2 significantly reduces the production of proinflammatory cytokines secreted by gastric adenocarcinoma (AGS) cells in response to *H. pylori* infection, suggesting that it may have potential for development as an anti-virulence drug that attenuates the pathogenicity of the pathogen (33).

Here, we describe the genetic and biochemical variability of antibiotic-resistant *H. pylori* strains isolated from patient biopsies in Québec (Canada) and two *H. pylori* control strains. Seven of the 13 analyzed strains carry the *cag*PAI. We characterized the CagA EPIYA and CM motifs as well as the amino-acid variations in the CagL protein showing considerable variability. We also analyzed the effects of co-cultivation with the *H. pylori* strains on IL-8 production, on cytoskeletal changes in epithelial gastric cells and on the production of extracellular T4SS pili. Finally, we demonstrated the effects of the 1G2 molecule on the strains suggesting that this molecule may have potential for development as an anti-virulence treatment.

## Materials and Methods

### Bacterial strains, cell lines and culture conditions

*H. pylori* strains 26695 (ATCC700392) and ATCC43504 were used as positive controls, and the previously described (24,33) Δ*cagV* (hp0530) mutant recreated in our laboratory was used as a negative control. *H. pylori* strains that were isolated from biopsies of 13 patients at the CHUM (Centre Hospitalier de l’Université de Montréal, Québec, Canada) (supplementary Table 1). All the strains were cultivated on Columbia agar base (BD) containing 10% (v/v) horse serum (Wisent Inc.), 5% (v/v) laked horse Blood (Wisent Inc.) with β-cyclodextrin (2 mg/ml), vancomycin (10 μg/ml) and amphotericin B (10 μg/ml). Chloramphenicol (34 μg/ml) was added in case of the Δ*cagV* strain to select for the chloramphenicol (*cam*) gene cassette used to disrupt the gene. For liquid cultures, brain heart infusion (BHI) media (Oxoid) was supplemented with 10% fetal bovine serum (FBS) and appropriate antibiotics. Bacteria were cultivated at 37°C under microaerophilic conditions (5% oxygen, 10% CO_2_). AGS cells (CRL-1739) were grown at 37°C in F12K media (Wisent Inc.) with 10% (v/v) FBS (Wisent Inc.) in a 5% CO_2_-containing atmosphere.

Antimicrobial susceptibility testing for *H. pylori* was done using metronidazole and clarithromycin Etest strips. *H. pylori* clinical strains and control strain, cultured on blood agar plates, were suspended in NaCl 0.45% 2 ml for a McFarland no. 2 standard, these suspensions were inoculated on 5% sheep blood agar plates with sterile swabs. The plates on which Etest strips were added, one strip by plate, were incubated at 37°C, in a microaerobic atmosphere (5% O_2_, 10% CO_2_, and 85% N_2_) for 72 h. The minimal inhibitory concentration (MIC) was the one at the intersection of *H. pylori* growth on the strip. The susceptible, intermediate and resistant criteria for clarithromycin were those of CLSI (Clinical and Laboratory Standards Institute) guidelines. The susceptible and resistant criteria for metronidazole were those of EUCAST (European Committee on Antimicrobial Susceptibility Testing) guidelines (34–36).

### Extraction of bacterial genomic DNA

The extraction of bacterial genomic DNA was performed using the GenElute Bacterial Genomic DNA kit (Sigma Aldrich), according to the instructions of the manufacturer.

### Infection of AGS cells with H. pylori strains

AGS cells cultivated at 7×10^5^ cells/well density in 6-well plates were infected for 4h with *H. pylori* at a multiplicity of infection (MOI) of 1:100. *H pylori* were first pre-incubated for 1h in F12K media with 10% FBS with or without 1G2 at a concentration of 200 µM at 37°C in a 5% CO_2_-containing atmosphere.

### Imaging the AGS cell “Hummingbird” phenotype by interference contrast microscopy

AGS cells cultivated at a density of 6×10^5^ cells/well in 6-well plates were infected with overnight cultures of *H. pylori* at a MOI of 100:1. After 4h of incubation under microaerophilic conditions, the media was removed, and the wells were washed twice with cold PBS. AGS cells were fixed with 2.5% glutaraldehyde and visualized with a Nikon Eclipse TE2000U microscope.

### Analysis of IL-8 secretion by AGS cells

After 4h of infection, supernatants were sampled and centrifuged at 15,000 *g* to remove cells and debris and frozen at -20°C. The level of IL-8 in cell culture supernatants was determined using a Human IL-8 Uncoated ELISA kit (Invitrogen, ThermoFisher Scientific Inc.).

### Sample preparation and western blotting

After 4h of infection, the cells were wash with phosphate-buffered saline (Wisent Inc.) and lysed in RIPA Buffer (50 mM Tris-HCl pH 8.0, 150 mM sodium chloride, 1.0% Igepal CA-630 NP-40, 0.5% sodium deoxycholate, 0.1% sodium dodecyl sulfate), complemented with a protease inhibitor cocktail for mammalian tissues (Sigma Aldrich) and a phosphatase inhibitor cocktail for tyrosine protein phosphatases, acid and alkaline phosphatases (Sigma Aldrich). The cells were harvested, incubated at 95°C for 5 min with SDS-PAGE sample buffer and centrifuged for 10 min at 10,000 rpm, followed by SDS-PAGE and western blotting. The production of proteins was assessed with monoclonal mouse anti-*Helicobacter pylori* CagA (HyTest Ltd.) and rabbit Cagα antiserum (Abcam). The actin has been identified using the Antibody Anti-β-Actin Antibody (C4): m-IgG Fc BP-HRP (Santa Cruz, sc-528515). The secondary antibodies (rabbit and mouse) were purchased from Biorad and the HRP signal was developed using Clarity Western ECL Substrate (Biorad).

### Sample preparation and scanning electron microscopic analysis

AGS cells were cultivated on round cover glasses (Fisherbrand) at 6×10^5^ cells/well density in 6-well plates. The cells were infected with *H pylori* for 4h as explained above, washed with cold phosphate buffer (PB 0.1 M), fixed in 4% paraformaldehyde/0.1% glutaraldehyde for 30 min at 4°C, followed by another wash in PB. The samples were then incubated in osmium tetroxide 4% (0.1%) for 1 hour at 4°C, followed by washing with PB. The samples were then dehydrated using a series of ethanol dilutions for 15 min each (30%, 50%, 70%, 80%, 90%, 95%, 100%, 100%), followed by drying using critical point dryer using the Leica EM CPD300. Then cells were then coated with 5 nm of carbon using the Leica EM ACE600. The sample were visualised using a Hitachi Regulus 8220 scanning electron Microscope. The length and number of T4SS pili was measured using ImageJ software.

### Statistical analysis

All the statistical analyses were performed using GraphPad Prism.

## Results

### Analysis of the cagPAI in H. pylori clinical isolates

We studied 13 *H. pylori* strains isolated from patient biopsies from 2017 to 2018 at the CHUM (Centre Hospitalier de l’Université de Montréal) presenting a resistance to clarithromycin and/or metronidazole, and two ATCC control strains, strain 26695 and ATCC43504 (supplementary Table 1). After extraction of bacterial genomic DNA, PCR amplification of the *cagA* and *cagL* genes was performed to determine the presence of the *cagPAI* (supplementary Figure 1, supplementary Table 2). The *vacA* gene, which encodes for the vacuolating cytotoxin and is present in virtually all *H. pylori* strains (37), was used as a control for PCR amplifications to confirm the identity of *H. pylori*; all strains were *vacA* positive (data not shown). Among the 13 clinical *H. pylori* strains, seven are *cagPAI-*positive as were the two reference strains from the ATCC (Table 1). Out of the seven *cagPAI-*positive strains, four (#3793, #3813, #3822 and #3830) are readily cultivatable under laboratory conditions and we selected them for the rest of this study.

**Table 1.**
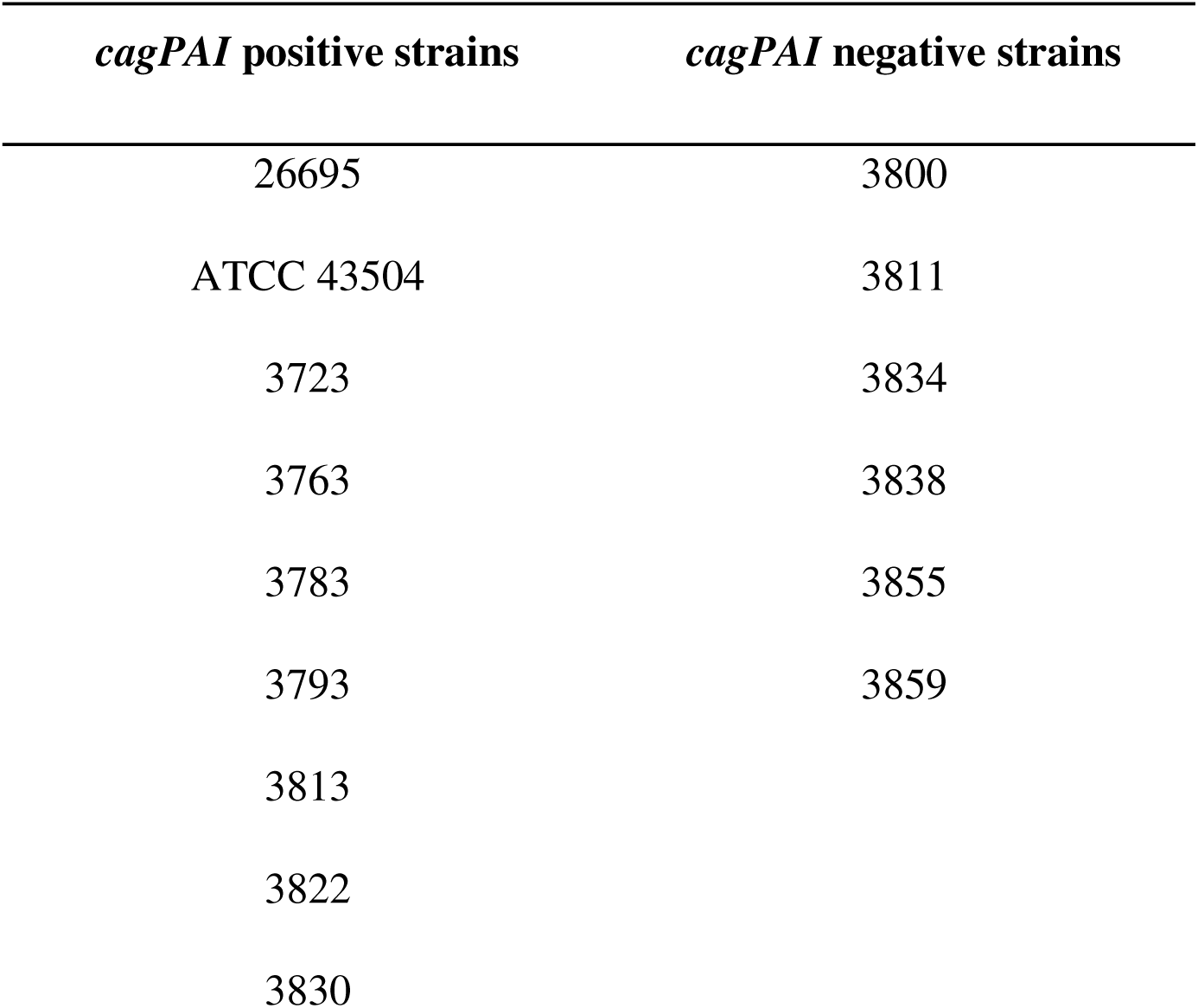
*cag*PAI positive and negative *H. pylori* strains.

| *cagPAI* positive strains | *cagPAI* negative strains |
| --- | --- |
| 26695 | 3800 |
| ATCC 43504 | 3811 |
| 3723 | 3834 |
| 3763 | 3838 |
| 3783 | 3855 |
| 3793 | 3859 |
| 3813 |  |
| 3822 |  |
| 3830 |  |

### Analysis of the variability of the CagA and CagL sequences

CagA displays C-terminal variability, which involves different types and/or numbers of EPIYA repeat sequences, and this diversity may correlate with differences of their virulence and distinct clinical outcomes (12,17). The *cagA* PCR products of the *cag*PAI positive strains were sequenced, followed by comparative analysis of the EPIYA motifs, CM types and A/T polymorphisms (Table 2 and supplementary Figures 2 and 3). Most of the strains comprise a Western CagA type (EPIYA-ABC) with two CM motifs, one inside the EPIYA-C and one distal to the EPIYA-C segment. Interestingly, the clinical isolate #3793 carries both the Western CM motif (CM^W^) and the East Asian CM motif (CM^EA^). The clinical strain #3830 contains an East Asian type (EPIYA-ABD) with one CM^EA^ motif distal to the EPIYA-C segment. Finally, the A/T polymorphism is present in the EPIYA-B motifs of the clinical strain #3822 and in the reference strain 26695.

**Table 2.**
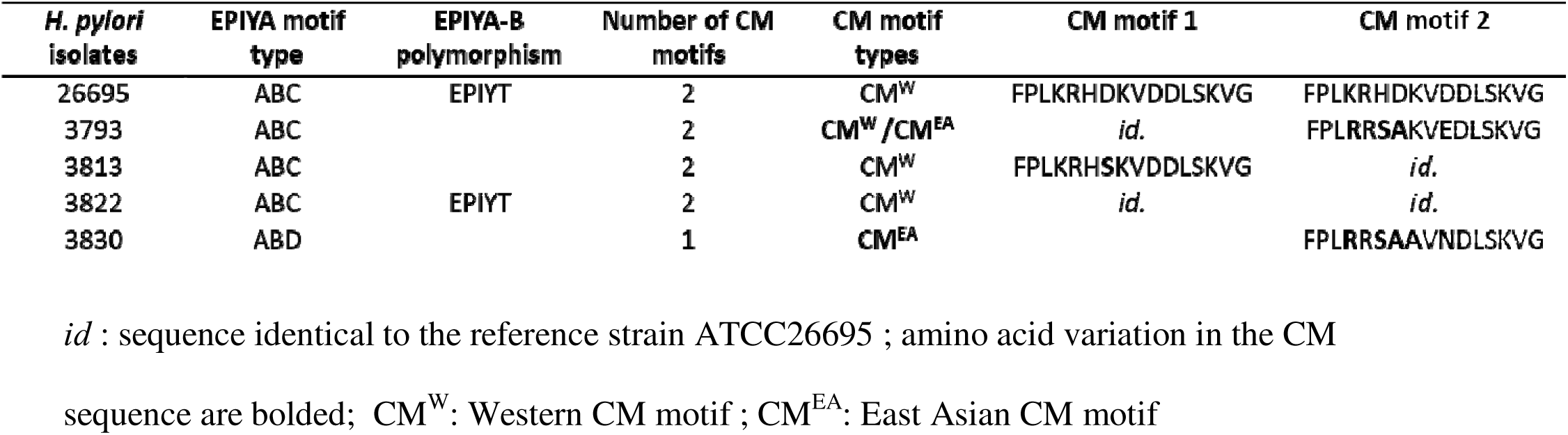
Characteristics of the CagA isoforms from the *cag*PAI positive *H. pylori* strains.

Next, we PCR-amplified the *cagL* gene, followed by sequencing and alignment to the CagL sequences from the *H. pylori* reference strain 26695 (Figure 1). The alignment shows that only two clinical strains (#3793 and #3822) do not contain the characteristic Glu59 residue, and that all of them contain the residues Asn201 and Ile234. These residues were previously reported to occur at higher rates in strains from gastric cancer patients (23). We also observe other sequence variations supporting that the strains have diverse origins.

**Figure 1.**
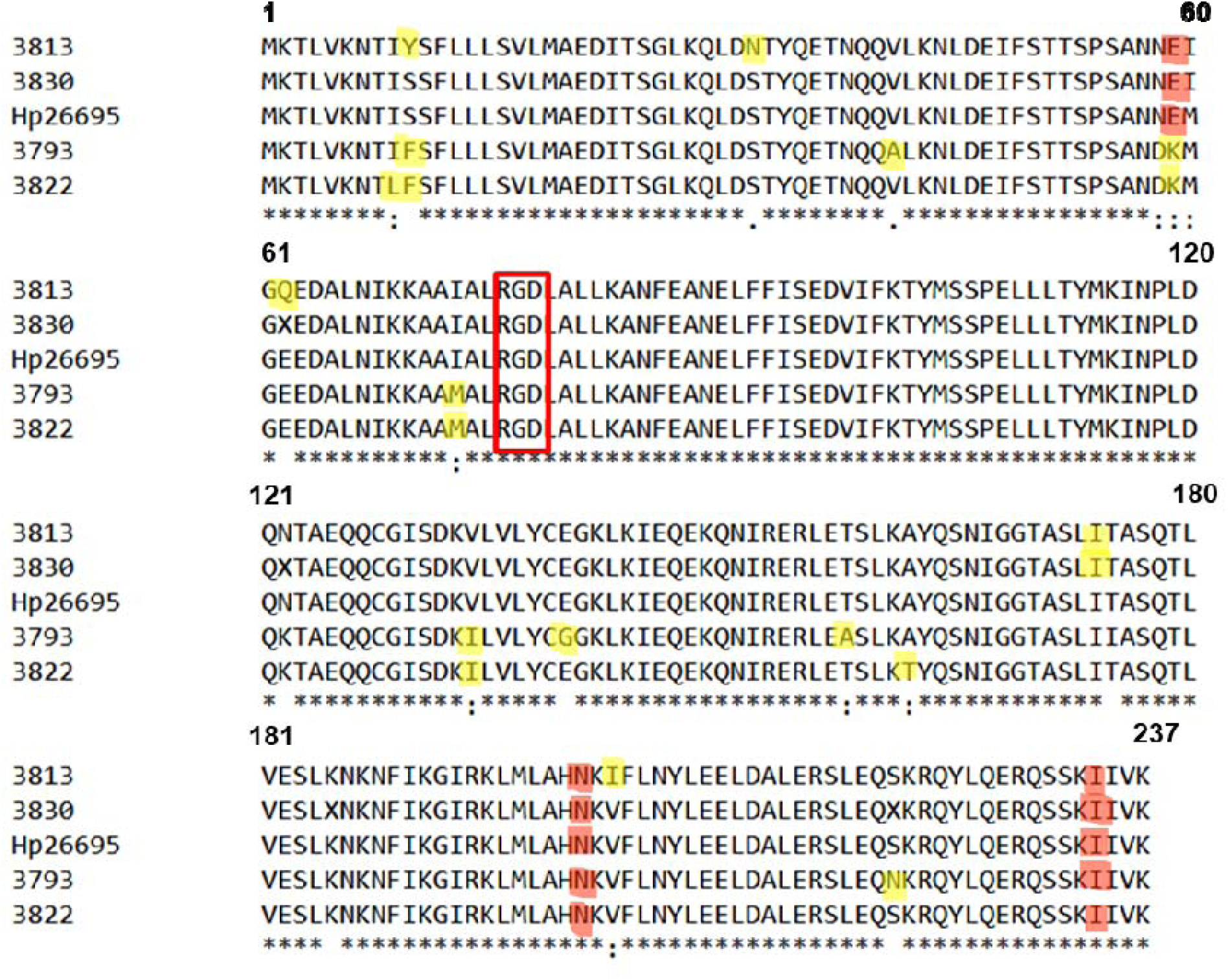
Alignment of the CagL sequences from the *cag*PAI positive *H. pylori* strains. Amino acid residues that occur at a significantly higher rate in isolates from patients with gastric cancer are highlighted in red, all the other variations identified within the alignment are highlighted in yellow. The highly conserved RGD motif that is involved in the interaction with host cell integrins is indicated by a red box.

### Effect of infection with H. pylori on co-cultivated AGS cells

CagA translocation into epithelial cells induces dysregulation of cellular signaling leading to the so-called “Hummingbird” phenotype, characterized by spreading and elongation of the cells (38). We co-cultivated AGS cells with the *H. pylori* strains, followed by interference contrast microscopy to monitor cell shape and counting of the proportion of cells displaying the Hummingbird phenotype (Figure 2). No morphological changes are observed in AGS cells alone and the number was low (up to 3%) in the case of cells incubated with the avirulent △*cagV* strain. Incubation with strains 26695, #3793 and #3813 induces the Hummingbird phenotype in 13% to 14% of the cells, respectively. In the case of strains #3822 and #3830, we observe that a larger number of cells (23% and 29%) display the characteristic morphological changes (Figure 2). We next analyzed the levels of IL-8 produced by the AGS cells in response to infection with the *H. pylori* strains. We observed that infection with the #3813 strains induces lower levels of IL-8 production than the 26695 reference strain. In contrast, strains #3793, #3822 and #3830 strains induce IL-8 production to comparable levels as 26695 (Figure 3).

**Figure 2.**
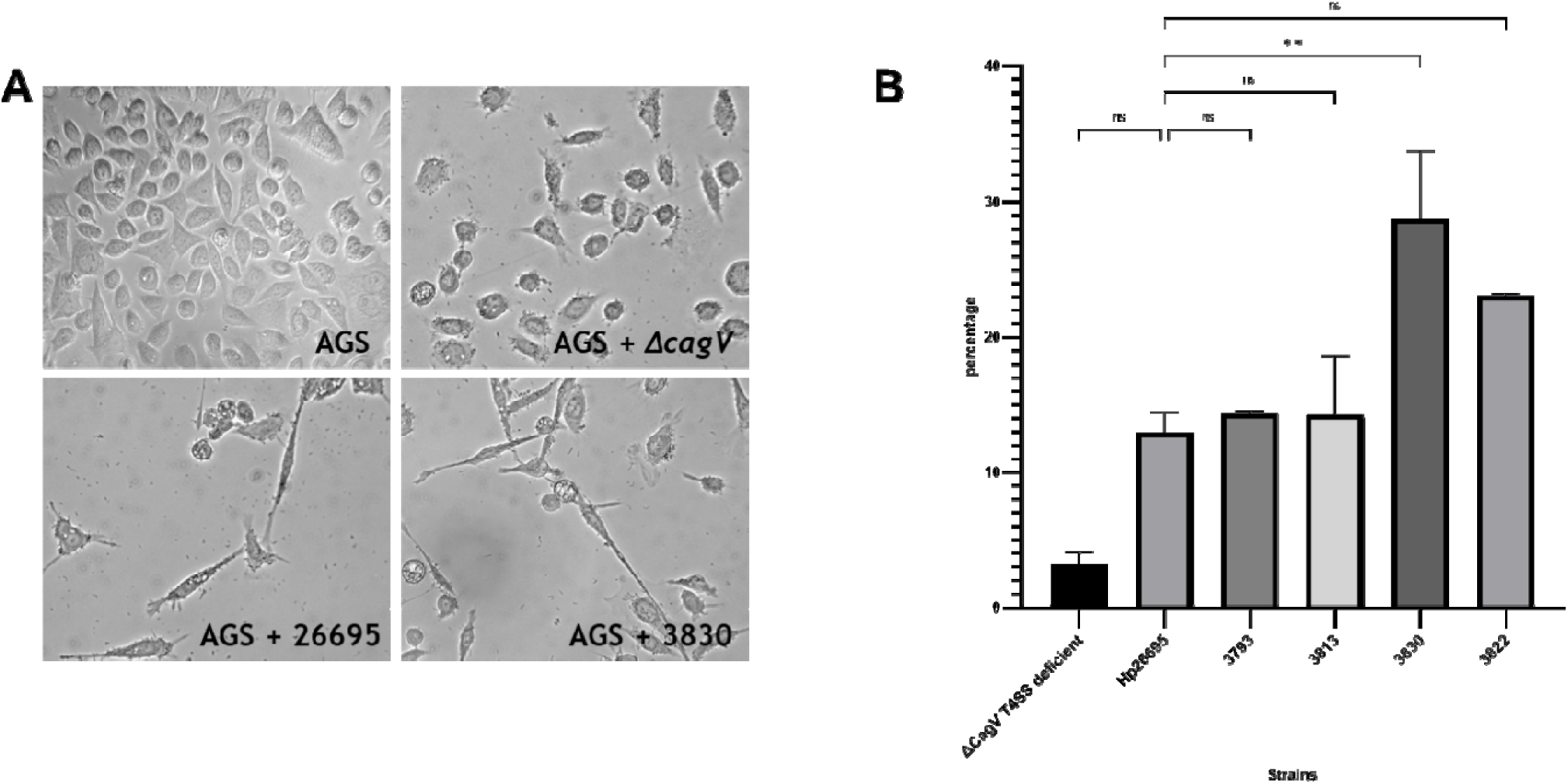
Hummingbird phenotype induced by *H. pylori* infection. (A) AGS cells were infected for four hours with *H. pylori* strains at MOI 100, followed by washing and the adherent cells were fixed and observed using interference contrast microscopy. Negative controls (AGS) and elongated cells induced by the infection with △*cagV,* Hp26695 and #3830 strains are shown as examples; each image represents a representative example for each condition. (B) Elongated cells were counted, and the results are represented as a percentage of total cells. The data represent means and standard deviations of duplicate experiments with at least 500 cells for each condition. **: p<0.05, Anova test.

**Figure 3.**
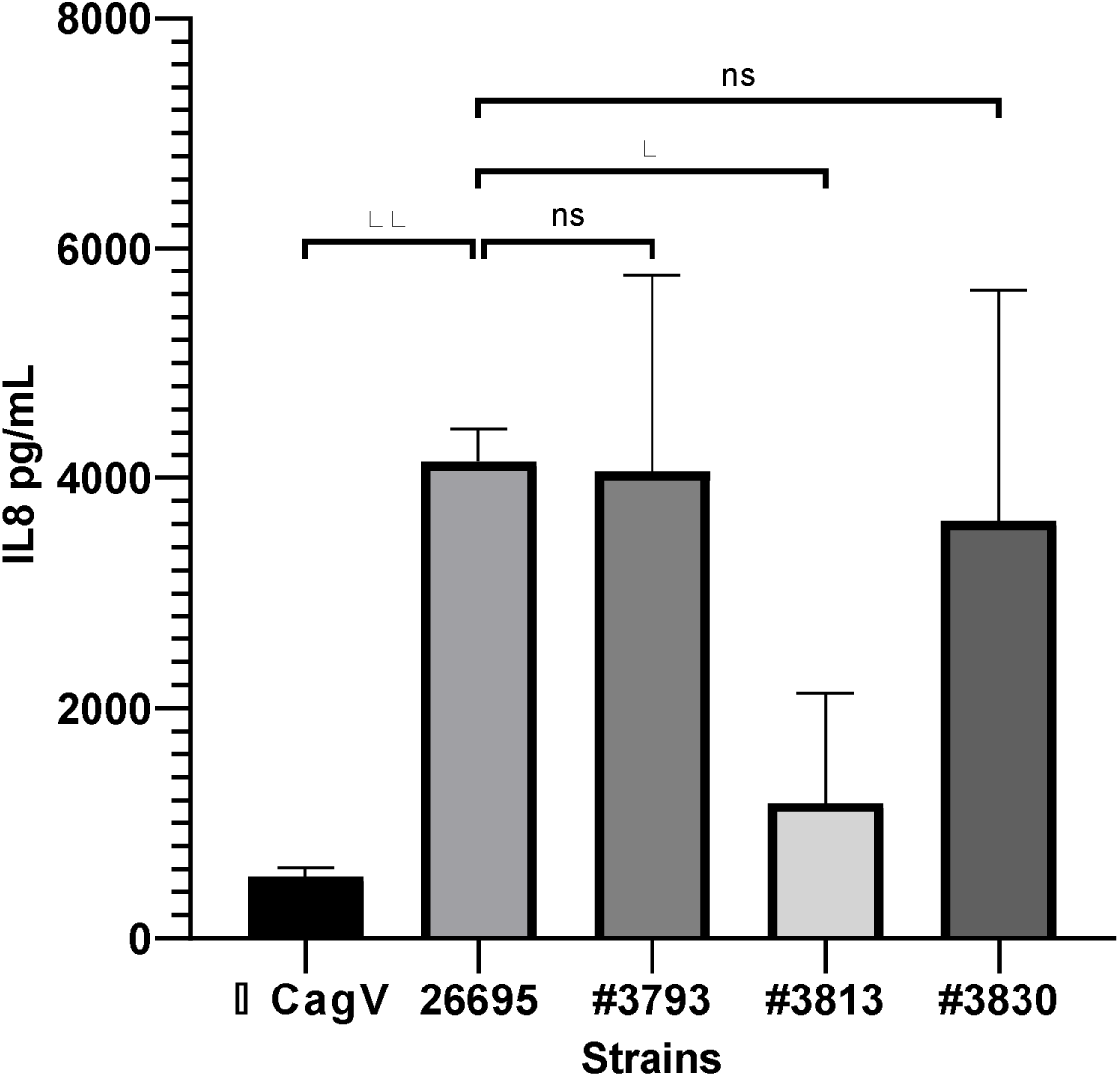
Interleukin-8 production by AGS cells infected with *H. pylori* strains. AGS cells were infected for 24h with the different *H. pylori* strains and the amounts of secreted IL-8 were measured by ELISA (n=4). ** p<0.005, * p< 0.01, ANOVA, ns: non-significant.

### Molecule 1G2 reduces IL-8 secretion of some H. pylori clinical isolates

We previously described the 1G2 molecule that inhibits Cagα ATPase activity and significantly reduces the pro-inflammatory response induced by infection with *H. pylori* strain 26695 (33). To assess whether this molecule has potential for application as inhibitor of the virulence of clinical strains we tested the impact on strains #3793, #3813, #3822 and #3830. We observed that co-incubation with 1G2 at 200 μM concentration significantly decreases the production IL-8 in the case of strain #3793 as well as in the 26695 control. In contrast, in the case of strains #3813, #3830 and #3822, we do not observe a significant difference of IL8 induction in the presence of 1G2 (Figure 4A). Since the inhibitor 1G2 may also affect the stability of its target, we also monitored levels of the oncoprotein CagA and of Cagα using western blotting (Figure 4B). We do not observe differences of the levels of CagA in the presence or in the absence of 1G2. In strain #3830 the protein is not detected by the antibody used in our experiment. Similarly, 1G2 has no effect on the levels of Cagα suggesting that it does not negatively impact the stability of Cag proteins.

**Figure 4.**
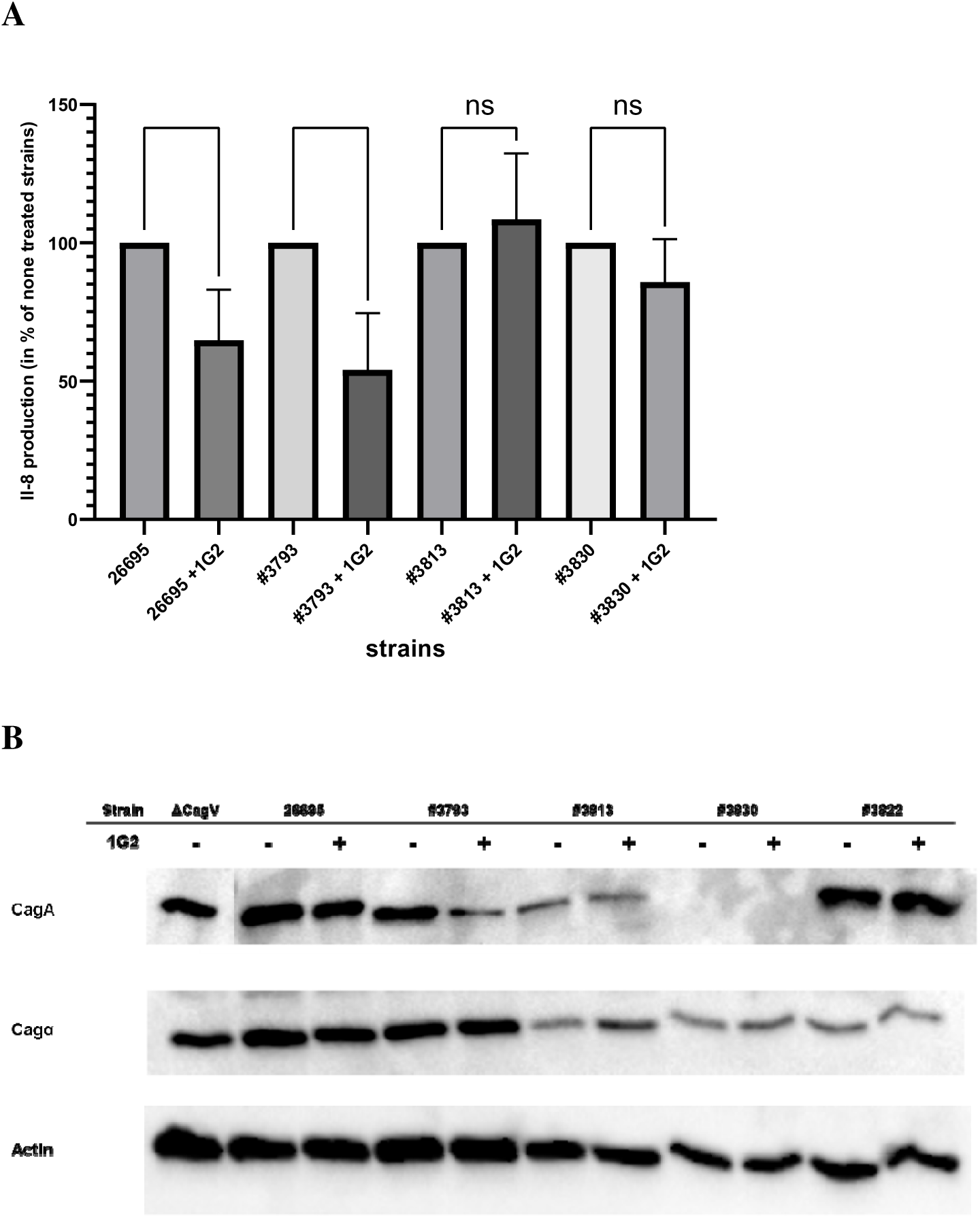
Effects of molecule 1G2 on IL-8 production on the levels of CagA and Cagα. *H. pylori* strains were pre-incubated without or with 200 μM of 1G2 for 1h. AGS cells were then co-cultured with *H. pylori* strains for 4h and IL-8 induction was measured by ELISA (A). The levels of CagA and Cagα were analyzed by SDS-PAGE of cell lysates, followed by western blotting with specific antisera; actin levels were also monitored as a loading control (B). The results of the IL-8 quantification were analyzed by pairs comparing the non-treated result (100% induction) with the treated results. The Δ*cagV* strain was used as negative control and analyzed using the strain 26695 as 100%. (n=6). ** p<0.005, * p< 0.01, ANOVA, ns: non-significant.

### Molecule 1G2 affects the formation of T4SS pili

As an alternative approach to measure the activity of the T4SS we monitored the formation of T4SS pili during the infection of AGS cells using scanning electron microscopy (SEM) in the presence or in the absence of 1G2 (Figure 5A). We observe an average of 14-15 T4SS pili per bacterium for the strains 26695 and #3793 in the absence of 1G2 (Figure 5B). In the presence of 1G2, the number of pili is drastically reduced, and we observe on average three T4SS pili per cell in both strains (Figure 5B). We do not observe any T4SS pili in strains #3813 and #3830 in the presence or in the absence of 1G2. Finally, in strain #3822 1G2 does not have an impact on the number of T4SS pili (12/cell), which correlates with the absence of an effect on IL8 induction (Figure 5 B, Figure 4A).

**Figure 5.**
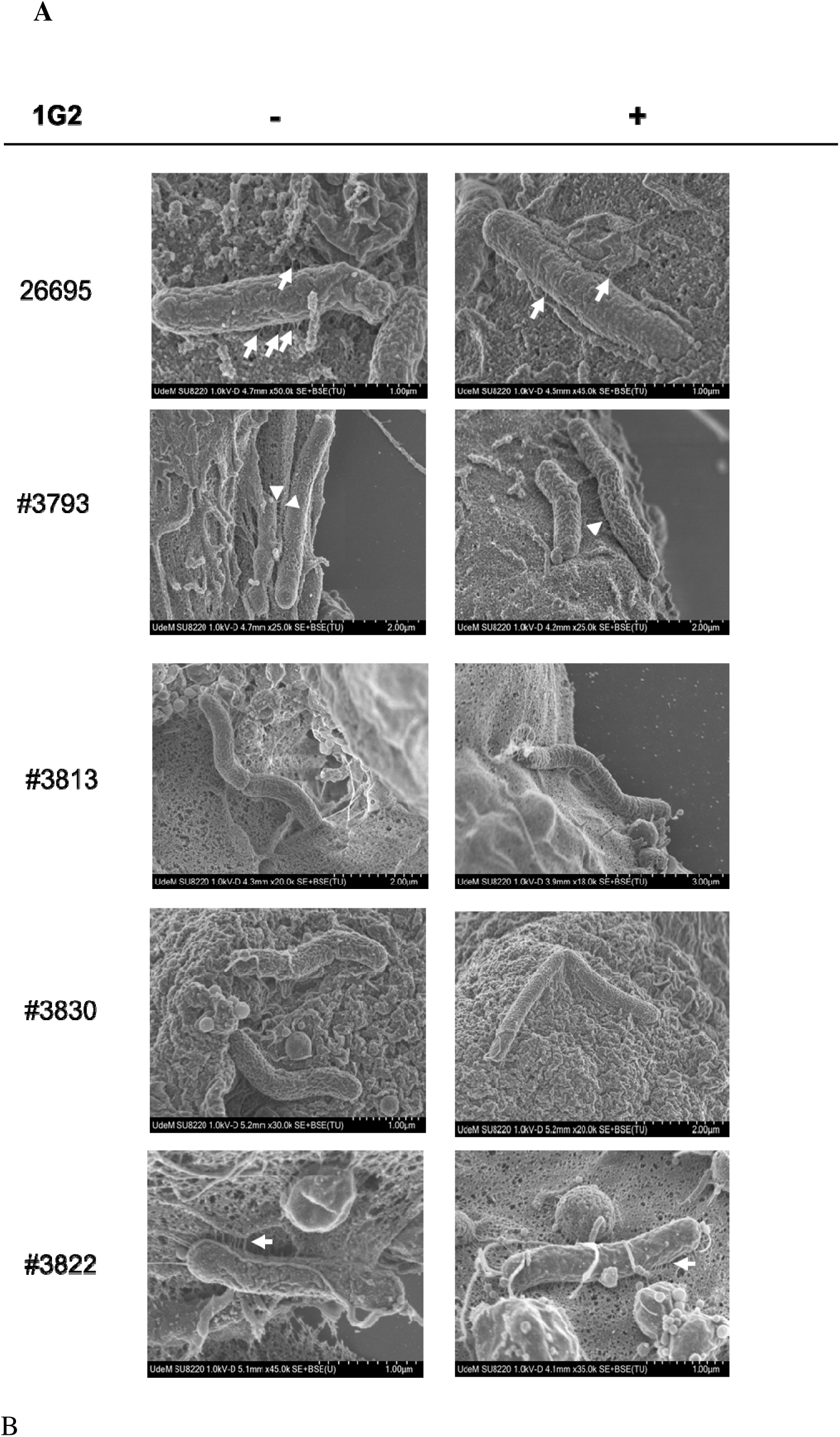
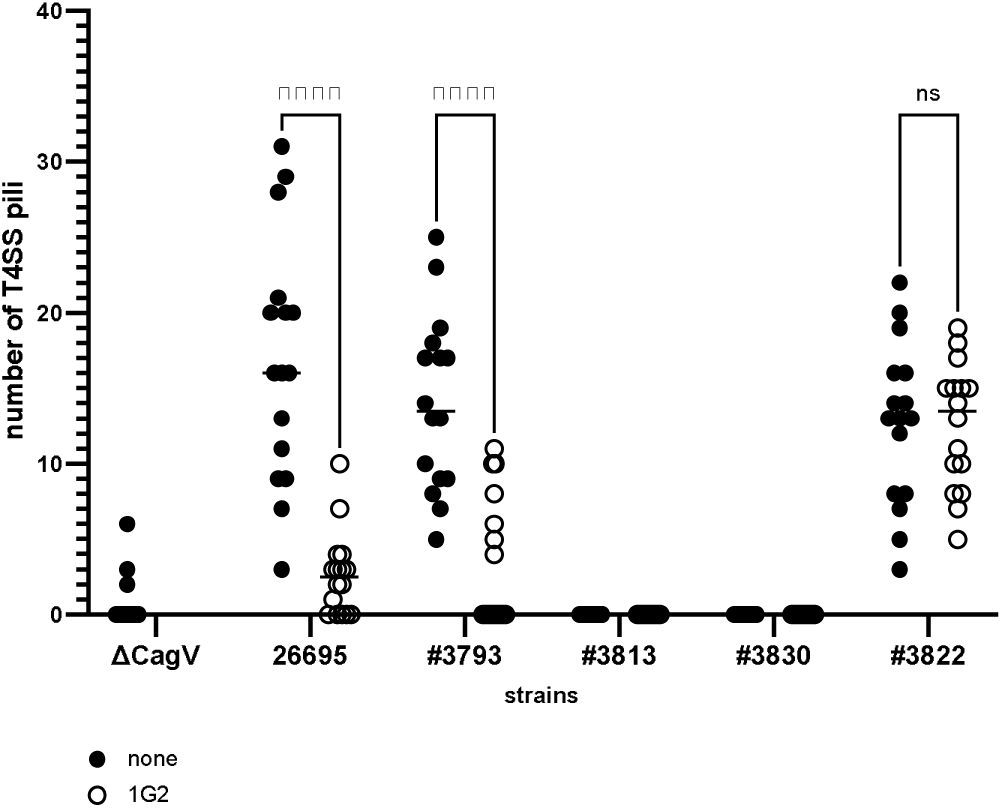
1G2 affects the number of T4SS pili during AGS infection. *H. pylori* strains were pre-incubated without (-) or with (+) 200 μM of 1G2, followed by co-cultivation with AGS cells for 4h and the samples were then fixed and analyzed using scanning electron microscopy. T4SS pili are identified with arrows (A) and the T4SSpili were counted using ImageJ (B) (n = 16 images). The Δ*cagV* strain was used as negative control. ****: p<0.001, Student’s T-test, ns: non-significant

## Discussion

The presence of a functional *cag* pathogenicity island (*cag*PAI) in *H. pylori* strains has been demonstrated to increase the risk for the development of more severe forms of gastric diseases (4–6,12). Here, we characterized 13 antibiotic-resistant *H. pylori* strains isolated from patient biopsies in Québec. We showed that seven of the strains carry the *cagA* and *cagL* genes and were thus considered as *cag*PAI positive and we further characterized the virulence phenotype of four strains (#3793, #3813, #3822 and #3830) that could be readily cultivated in laboratory conditions. To assess the virulence of *H. pylori* strains we monitored the production of Cag proteins, the induction of IL-8 production, the induction of changes of the cytoskeleton (“hummingbird” phenotype) and the formation of T4SS pili. The clinical strains induce a hummingbird phenotype in AGS cells that is similar to the 26695 control and the cytoskeletal changes are even more pronounced in strain #3830. Similarly, all clinical strains induce IL-8 production in AGS cells. These data suggest that all the clinical strains carry functional T4SSs and that their effects can be monitored using the established AGS cell infection assay.

We also monitored the production of T4SS pili after co-incubation with AGS cells by scanning electron microscopy and this constitutes a novel quantifiable assay for T4SS function. We observe between 12 and 15 T4SS pili per cell in the 26995 strain and in the clinical strains #3793 and #3822. In contrast, we do not observe any T4SS pili in strains #3813, #3830. Both strains induce IL-8 production and cytoskeletal changes after co-incubation with AGS cells suggesting that they do not need elaborate extracellular structures to translocate virulence factors. This constitutes a marked difference to previously characterized *H. pylori* strains even if we cannot exclude that these strains assemble very short T4SS pili that can not be detected by SEM. In addition, the imaging was done using only one angle, limiting the observation of the interaction between the bacteria and cells.

Comparative analysis of the CagA C-terminal domains showed significant variability among the *H. pylori* strains. Interestingly, strain #3793 carries both eastern and western CM motifs (CM^W^/CM^EA^). The CM motif has been demonstrated to serve as a binding site for PAR1b and this interaction is highly influenced by the nature of the motif and its numbers of repetition (39). The presence of this combination of CM motifs may therefore influence the virulence of strain #3793, but differences are not apparent when IL-8 production or effects on cytoskeletal changes are monitored. Similarly, the differences between the CagL sequences of the different strain do not correlate with effects on AGS cells, but they may impact the pathogenicity in patients.

Finally, we tested the effects of the previously characterized 1G2 molecule on the antibiotic resistant *H. pylori* strains (33). We showed that 1G2 significantly decreases T4SS activity in strains 26695 and #3793 as measured by IL-8 production of AGS cells and assembly of T4SS pili. In contrast, 1G2 has no effect on the production of IL-8 in strains #3813, #3830 and #3822 or on the production of T4SS pili in strain #3822. These results suggest that 1G2 may have potential for the development of treatments against some clinical strains, but that more potent derivatives would have to be developed to have broad clinical impact. 1G2 has no effect on the levels of CagA or of the target Cagα showing that it does not destabilize its target. Interestingly, 1G2 inhibits the formation of T4SS pili in strains 26695 and #3793 suggesting that this is the primary mechanism for the effect of this molecule on *H. pylori* virulence.

## Acknowledgments

This work was supported by grants from the Cancer Research Society, the Charles Bowers Memorial Fund, the Bergeron-Jetté Foundation (CRS, #23404 and #25102) and the Natural Sciences and Engineering Research Council (NSERC, #RGPIN-2017-05123) to C.B. We grateful to Dr. Dainelys Guadarrama Bell and Dr. Nanci at the Université de Montréal electron microscopy facility for technical support and assistance.

## Author contributions

F.O. designed and carried out experiments and analyzed data; C.M. designed and carried out experiments and analyzed data; M.B. contributed resources, analyzed data; C.G. contributed resources, analyzed data; C.B. designed experiments, analyzed data and wrote the manuscript with input from all the co-authors. All co-authors have approved the final version of this manuscript.

## Study materials

Data, analytic methods, and study materials will be made available to other researchers upon request addressed to the corresponding author.

## Declaration of interests

The authors declare no competing interests.

## Supplementary data

**Supplementary table 1.** *H. pylori* strains and resistance to antibiotics.

| *H. pylori* Strain | Resistance |  |
| --- | --- | --- |
|  | Metronidazole (MIC mg/L) | Clarithromycin (MIC mg/L) |
| 3723 | R (256) | R (256) |
| 3763 | R (> 256) | R (> 256) |
| 3783 | S (0.06) | R (8) |
| 3793 | R (> 256) | R (256) |
| 3800 | R (128) | S (<= 0.03) |
| 3811 | R (> 256) | R (1) |
| 3813 | R (128) | R (256) |
| 3822 | R (> 256) | R(> 256) |
| 3830 | S (0.06) | S (<= 0.03) |
| 3834 | R (> 256) | R (32) |
| 3838 | R (> 256) | R (16) |
| 3855 | R (> 256) | R (32) |
| 3859 | R (64) | S (<= 0.03) |
| ATCC 43504 | R | S |
| 26695 | S | S |

S : susceptible, R : resistant

**Supplementary table 2.**
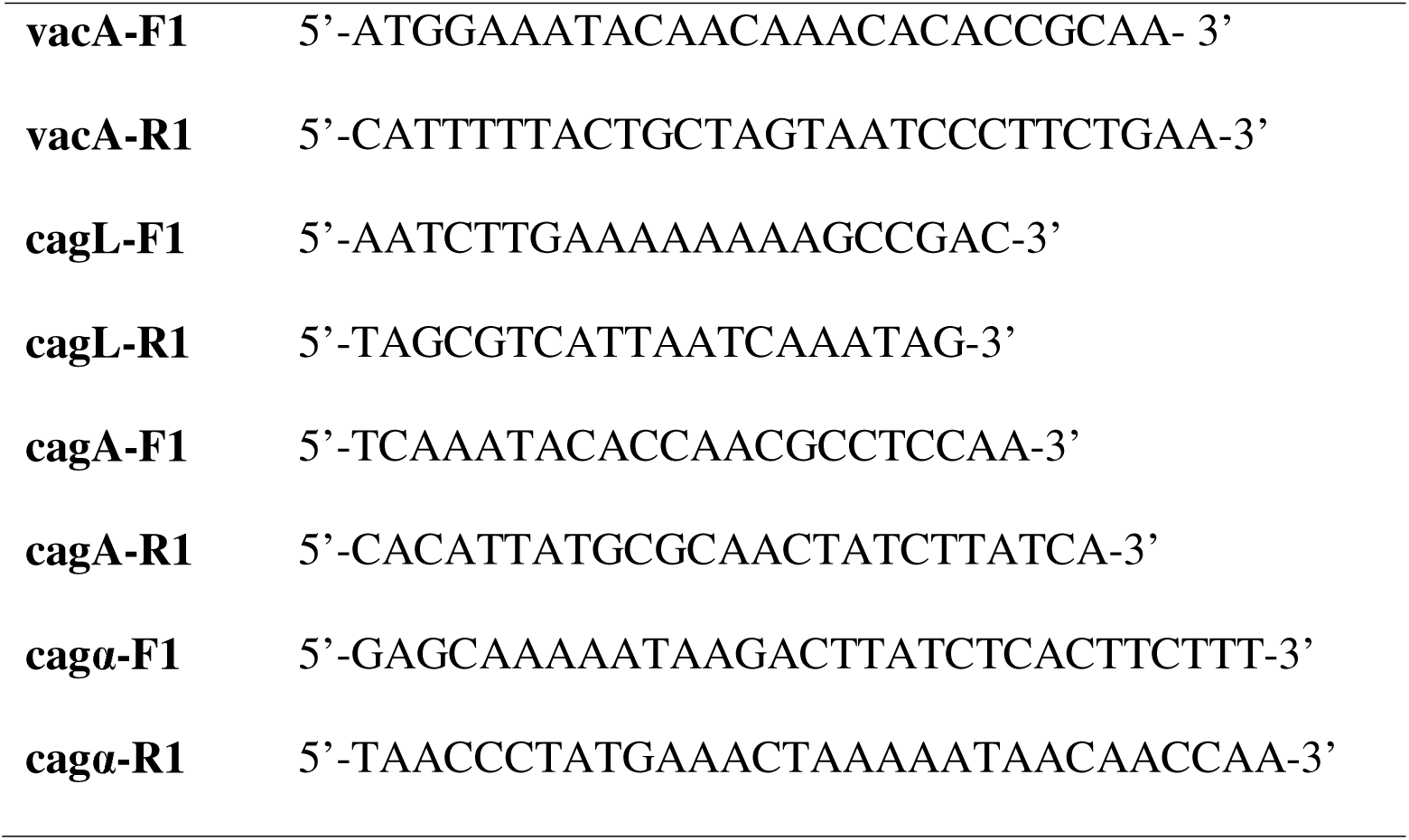
Primers used for PCR amplification and sequencing.

**Supplementary Figure 1.**
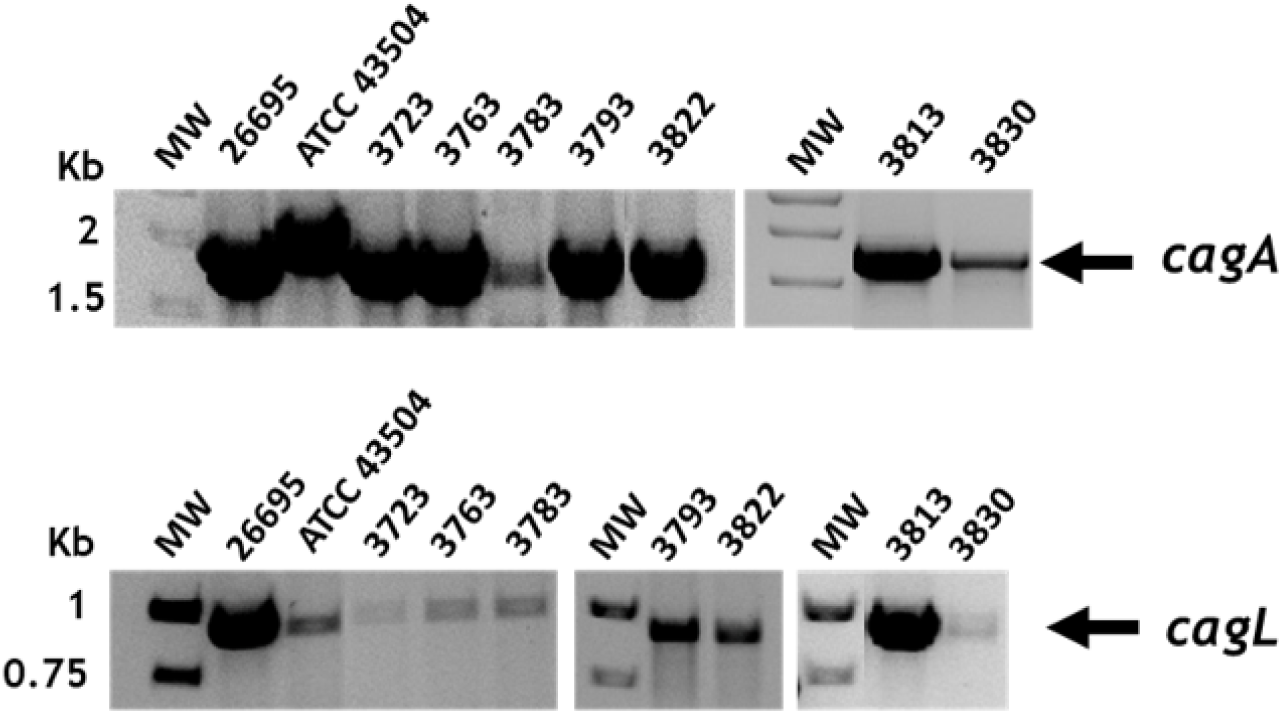
PCR amplification of *cagA* and *cagL* genes from *H. pylori* isolates. PCR amplifications with *cagA*- and *cagL*-specific primers were performed from genomic DNA isolated from the *H. pylori* strains followed by agarose gel analysis. Kb: Kilobases.

**Supplementary Figure 2.**
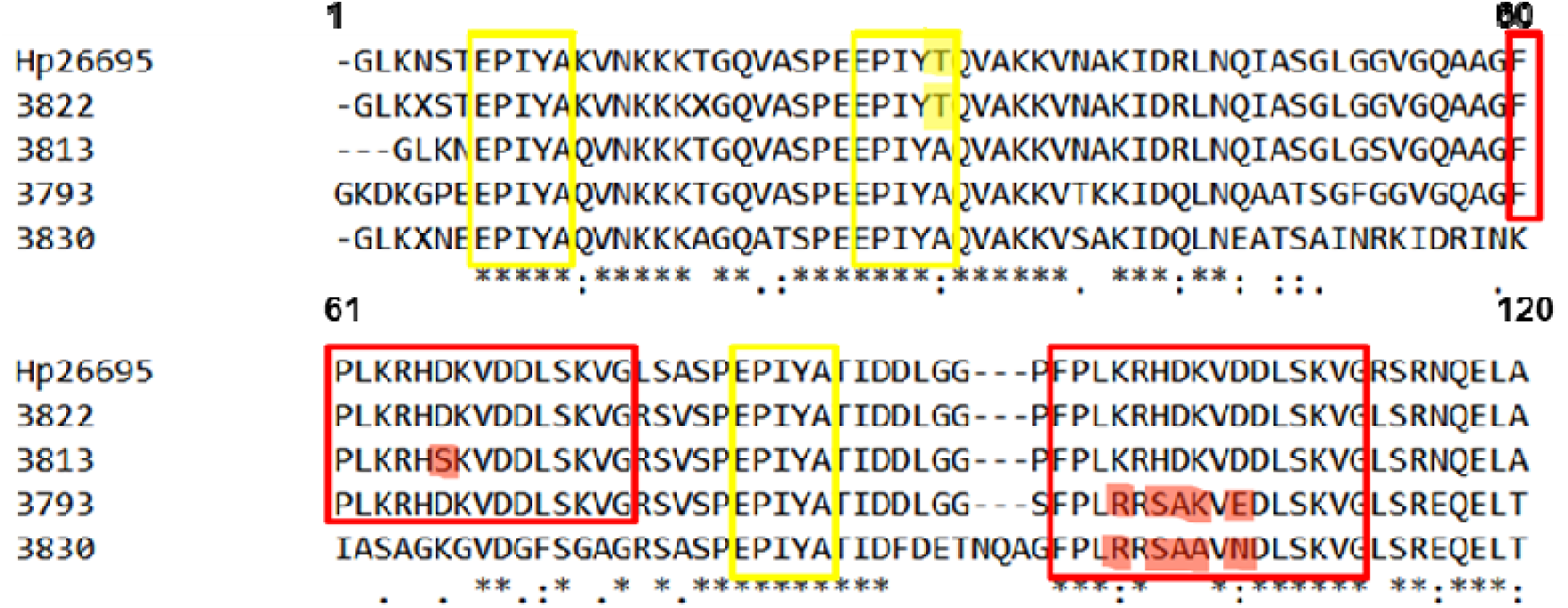
Alignment of the sequences of CagA comprising the three EPIYA motifs. The EPIYA motifs are labelled with yellow boxes and the CM motifs are marked with red boxes; variations inside the sequence are highlighted in the same colors.

## Notes

### Competing Interest Statement

The authors have declared no competing interest.

## References

1. Covacci A, Telford JL, Giudice GD, Parsonnet J, Rappuoli R. *Helicobacter pylori* Virulence and Genetic Geography. Science. 1999 May 21;284(15418):1328–33.

2. Cover TL, Blaser MJ. Helicobacter pylori factors associated with disease. Gastroenterology. 1999 Jul;117(1):257–60.

3. Gatta L, Vakil N, Vaira D, Scarpignato C. Global eradication rates for Helicobacter pylori infection: systematic review and meta-analysis of sequential therapy. BMJ. 2013 Aug 7;347(aug07 1):f4587–f4587.

4. Cover TL. *Helicobacter pylori* Diversity and Gastric Cancer Risk. mBio. 2016 Mar 2;7(1):e01869–15, /mbio/7/1/e01869-15.atom.

5. Yong X, Tang B, Li BS, Xie R, Hu CJ, Luo G, et al. Helicobacter pylori virulence factor CagA promotes tumorigenesis of gastric cancer via multiple signaling pathways. Cell Commun Signal. 2015 Dec;13(1):30.

6. Hatakeyama M. Helicobacter pylori CagA and Gastric Cancer: A Paradigm for Hit-and-Run Carcinogenesis. Cell Host & Microbe. 2014 Mar;15(3):306–16.

7. Odenbreit S. Translocation of Helicobacter pylori CagA into Gastric Epithelial Cells by Type IV Secretion. Science. 2000 Feb 25;287(5457):1497–500.

8. Viala J, Chaput C, Boneca IG, Cardona A, Girardin SE, Moran AP, et al. Nod1 responds to peptidoglycan delivered by the Helicobacter pylori cag pathogenicity island. Nat Immunol. 2004 Nov;5(11):1166–74.

9. Varga MG, Shaffer CL, Sierra JC, Suarez G, Piazuelo MB, Whitaker ME, et al. Pathogenic Helicobacter pylori strains translocate DNA and activate TLR9 via the cancer-associated cag type IV secretion system. Oncogene. 2016 Dec;35(48):6262–9.

10. Gall A, Gaudet RG, Gray-Owen SD, Salama NR. TIFA Signaling in Gastric Epithelial Cells Initiates the *cag* Type 4 Secretion System-Dependent Innate Immune Response to *Helicobacter pylori* Infection. Blaser MJ, editor. mBio. 2017 Sep 6;8(4):e01168–17, /mbio/8/4/e01168-17.atom.

11. Tegtmeyer N, Neddermann M, Asche CI, Backert S. Subversion of host kinases: a key network in cellular signaling hijacked by *Helicobacter pylori* CagA: Host kinases hijacked by CagA. Molecular Microbiology. 2017 Aug;105(3):358–72.

12. Hatakeyama M, Higashi H. Helicobacter pylori CagA: a new paradigm for bacterial carcinogenesis. Cancer Science. 2005 Dec;96(12):835–43.

13. Backert S, Moese S, Selbach M, Brinkmann V, Meyer TF. Phosphorylation of tyrosine 972 of the Helicobacter pylori CagA protein is essential for induction of a scattering phenotype in gastric epithelial cells: Function of the H. pylori CagA protein. Molecular Microbiology. 2008 Jul 7;42(3):631–44.

14. Tammer I, Brandt S, Hartig R, König W, Backert S. Activation of Abl by Helicobacter pylori: A Novel Kinase for CagA and Crucial Mediator of Host Cell Scattering. Gastroenterology. 2007 Apr;132(4):1309–19.

15. Selbach M, Moese S, Hauck CR, Meyer TF, Backert S. Src Is the Kinase of the Helicobacter pylori CagA Protein in Vitro and in Vivo. Journal of Biological Chemistry. 2002 Mar;277(9):6775–8.

16. Higashi H, Tsutsumi R, Fujita A, Yamazaki S, Asaka M, Azuma T, et al. Biological activity of the Helicobacter pylori virulence factor CagA is determined by variation in the tyrosine phosphorylation sites. Proceedings of the National Academy of Sciences. 2002 Oct 29;99(22):14428–33.

17. Zhang XS, Tegtmeyer N, Traube L, Jindal S, Perez-Perez G, Sticht H, et al. A Specific A/T Polymorphism in Western Tyrosine Phosphorylation B-Motifs Regulates Helicobacter pylori CagA Epithelial Cell Interactions. Blanke SR, editor. PLoS Pathog. 2015 Feb 3;11(2):e1004621.

18. Ren S, Higashi H, Lu H, Azuma T, Hatakeyama M. Structural Basis and Functional Consequence of Helicobacter pylori CagA Multimerization in Cells. Journal of Biological Chemistry. 2006 Oct;281(43):32344–52.

19. Wiedemann T, Hofbaur S, Tegtmeyer N, Huber S, Sewald N, Wessler S, et al. *Helicobacter pylori* CagL dependent induction of gastrin expression via a novel αvβ _5_ -integrin–integrin linked kinase signalling complex. Gut. 2012 Jul;61(7):986–96.

20. Barden S, Niemann HH. Adhesion of Several Cell Lines to Helicobacter pylori CagL Is Mediated by Integrin αVβ6 via an RGDLXXL Motif. Journal of Molecular Biology. 2015 Mar;427(6):1304– 15.

21. Kwok T, Zabler D, Urman S, Rohde M, Hartig R, Wessler S, et al. Helicobacter exploits integrin for type IV secretion and kinase activation. Nature. 2007 Oct;449(7164):862–6.

22. Yeh YC, Chang WL, Yang HB, Cheng HC, Wu JJ, Sheu BS. *H. pylori cagL* amino acid sequence polymorphism Y58E59 induces a corpus shift of gastric integrin α5β1 related with gastric carcinogenesis: *H. pylori* cagL & GASTRIC INTEGRIN. Mol Carcinog. 2011 Oct;50(10):751–9.

23. Ogawa H, Iwamoto A, Tanahashi T, Okada R, Yamamoto K, Nishiumi S, et al. Genetic variants of Helicobacter pylori type IV secretion system components CagL and CagI and their association with clinical outcomes. Gut Pathog. 2017 Dec;9(1):21.

24. Fischer W, Püls J, Buhrdorf R, Gebert B, Odenbreit S, Haas R. Systematic mutagenesis of the Helicobacter pylori cag pathogenicity island: essential genes for CagA translocation in host cells and induction of interleukin-8: Functional dissection of the H. pylori type IV secretion system. Molecular Microbiology. 2002 Jan 13;42(5):1337–48.

25. Jurik A, Haußer E, Kutter S, Pattis I, Praßl S, Weiss E, et al. The Coupling Protein Cagβ and Its Interaction Partner CagZ Are Required for Type IV Secretion of the *Helicobacter pylori* CagA Protein. Infect Immun. 2010 Dec;78(12):5244–51.

26. Kutter S, Buhrdorf R, Haas J, Schneider-Brachert W, Haas R, Fischer W. Protein Subassemblies of the *Helicobacter pylori* Cag Type IV Secretion System Revealed by Localization and Interaction Studies. J Bacteriol. 2008 Mar 15;190(6):2161–71.

27. Hilleringmann M, Pansegrau W, Doyle M, Kaufman S, MacKichan ML, Gianfaldoni C, et al. Inhibitors of Helicobacter pylori ATPase Cagα block CagA transport and cag virulence. Microbiology. 2006 Oct 1;152(10):2919–30.

28. Krause S, Bárcena M, Pansegrau W, Lurz R, Carazo JM, Lanka E. Sequence-related protein export NTPases encoded by the conjugative transfer region of RP4 and by the *cag* pathogenicity island of *Helicobacter pylori* share similar hexameric ring structures. Proc Natl Acad Sci USA. 2000 Mar 28;97(7):3067–72.

29. Savvides SN. VirB11 ATPases are dynamic hexameric assemblies: new insights into bacterial type IV secretion. The EMBO Journal. 2003 May 1;22(9):1969–80.

30. Yeo HJ, Savvides SN, Herr AB, Lanka E, Waksman G. Crystal Structure of the Hexameric Traffic ATPase of the Helicobacter pylori Type IV Secretion System. Molecular Cell. 2000 Dec;6(6):1461–72.

31. Machón C, Rivas S, Albert A, Goñi FM, de la Cruz F. TrwD, the Hexameric Traffic ATPase Encoded by Plasmid R388, Induces Membrane Destabilization and Hemifusion of Lipid Vesicles. J Bacteriol. 2002 Mar 15;184(6):1661–8.

32. Hare S, Fischer W, Williams R, Terradot L, Bayliss R, Haas R, et al. Identification, structure and mode of action of a new regulator of the Helicobacter pylori HP0525 ATPase. EMBO J. 2007 Nov 28;26(23):4926–34.

33. Arya T, Oudouhou F, Casu B, Bessette B, Sygusch J, Baron C. Fragment-based screening identifies inhibitors of ATPase activity and of hexamer formation of Cagα from the Helicobacter pylori type IV secretion system. Sci Rep. 2019 Dec;9(1):6474.

34. Wayne P. Methods for Antimicrobial Dilution and Disk Susceptibility Testing of Infrequently Isolated or Fastidious Bacteria [Internet]. 2016. 3rd p. (Clinical and Laboratory Standards Institute). Available from: https://webstore.ansi.org/preview-pages/CLSI/preview_CLSI+M45-Ed3.pdf

35. Best LM, Haldane DJM, Keelan M, Taylor DE, Thomson ABR, Loo V, et al. Multilaboratory Comparison of Proficiencies in Susceptibility Testing of *Helicobacter pylori* and Correlation between Agar Dilution and E Test Methods. Antimicrob Agents Chemother. 2003 Oct;47(10):3138–44.

36. eucast: Clinical breakpoints and dosing of antibiotics [Internet]. [cited 2023 Jul 19]. Available from: https://www.eucast.org/clinical_breakpoints

37. Cover TL, Blanke SR. Helicobacter pylori VacA, a paradigm for toxin multifunctionality. Nat Rev Microbiol. 2005 Apr;3(4):320–32.

38. Segal ED, Cha J, Lo J, Falkow S, Tompkins LS. Altered states: Involvement of phosphorylated CagA in the induction of host cellular growth changes by *Helicobacter pylori*. Proc Natl Acad Sci USA. 1999 Dec 7;96(25):14559–64.

39. Nishikawa H, Hayashi T, Arisaka F, Senda T, Hatakeyama M. Impact of structural polymorphism for the Helicobacter pylori CagA oncoprotein on binding to polarity-regulating kinase PAR1b. Sci Rep. 2016 Sep;6(1):30031.

